# Linked machine learning classifiers improve species classification of fungi when using error-prone long-reads on extended metabarcodes

**DOI:** 10.1101/2021.05.01.442223

**Authors:** Tavish Eenjes, Yiheng Hu, Laszlo Irinyi, Minh Thuy Vi Hoang, Leon M. Smith, Celeste C. Linde, Andrew W. Milgate, Wieland Meyer, Eric A. Stone, John P. Rathjen, Benjamin Mashford, Benjamin Schwessinger

**Affiliations:** Research School of Biology, Australian National University, Acton, ACT, 2601, Australia; Molecular Mycology Research Laboratory, Centre for Infectious Diseases and Microbiology, Faculty of Medicine and Health, Sydney Medical School, Westmead Clinical School, The University of Sydney, Camperdown, NSW, 2006, Australia; Sydney Institute for Infectious Diseases, The University of Sydney, Camperdown, NSW, 2006, Australia; Westmead Institute for Medical Research, Westmead, NSW, 2145, Australia; Wagga Wagga Agricultural Institute, New South Wales Department of Primary Industries, Wagga Wagga NSW 2650, Australia; Westmead Hospital (Research and Education Network), Westmead, NSW, 2145, Australia; Curtin Medical School, Curtin University, Perth Bentley, WA 6102, Australia; Biological Data Science Institute, Australian National University, Acton, ACT, 2601, Australia; Department of Microbial Interactions, IMIT/ZMBP, University of Tübingen, Tübingen, 72074, Germany; Diversity Arrays Technology, Bruce, ACT, 2617, Australia

**Keywords:** Machine learning, fungi, long-read sequencing, MinION, convolutional neural network

## Abstract

**Background:** The increased usage of error-prone long-read sequencing for metabarcoding of fungi has not been matched with adequate public databases and concomitant analysis approaches. We address this gap and present a proof-of-concept study for classifying fungal taxa using linked machine learning classifiers. We demonstrate the capability of linked machine learning classifiers to accurately classify species and strains using real-world and simulated fungal ribosomal DNA datasets, including plant and human pathogens. We benchmark our new approach in comparison to current alignment and k-mer based methods based on synthetic mock communities. We also assess real world applications of species identification in complex unlabelled datasets.

**Results:** Our machine learning approach assigned individual nanopore long-read amplicon sequences to fungal species with high recall rates and low false positive rates. Importantly, our approach successfully distinguished between closely-related species and strains when individual read errors were higher than the genetic distance between individual taxa, which the alignment and k-mer methods could not do. The machine learning approach showed an ability to identify key species with high recall rates, even in complex samples of unknown species composition.

**Conclusions:** A proof of concept machine learning approach using a tree-descent approach on a decision tree of classifiers can identify known taxa with high accuracy, and precisely detect known target species from complex samples with high recall rates. We propose this approach is suitable for detecting the known knowns of pathogens or invasive species in any environment of mostly unknown composition, including agriculture and wild ecosystems.

## BACKGROUND

DNA sequencing is increasingly becoming an important part of detecting, identifying and classifying fungal species, particularly through DNA barcoding [1]. To date, this process mostly involves the use of short, variable regions of DNA that differ between species and are surrounded by conserved regions suitable for ‘universal’ primers, enabling PCR amplification over a large variety of fungal taxa [2, 3]. The internal transcribed spacer (ITS) region is used as the primary DNA barcode region for fungal diversity studies [4]. This region contains the two variable components, ITS1 and ITS2, which are a combined 550-600 bp long on average [5]. ITS1 and ITS2 are separated by the conserved 5.8S rRNA gene and flanked by the conserved 18S and 28S rRNA genes (Figure 1a). Although these regions offer a targetable region for identifying fungal species, they have limitations that affect the ability to accurately classify fungi especially at lower taxonomic ranks [5, 6]. The length of the complete ITS1/2 region prevents short-read sequencing platforms to use both in combination for taxonomic classification. Furthermore, the limited selection of ‘universal’ primers in the region can subject taxonomic studies to primer biases [7].

**Figure 1:**
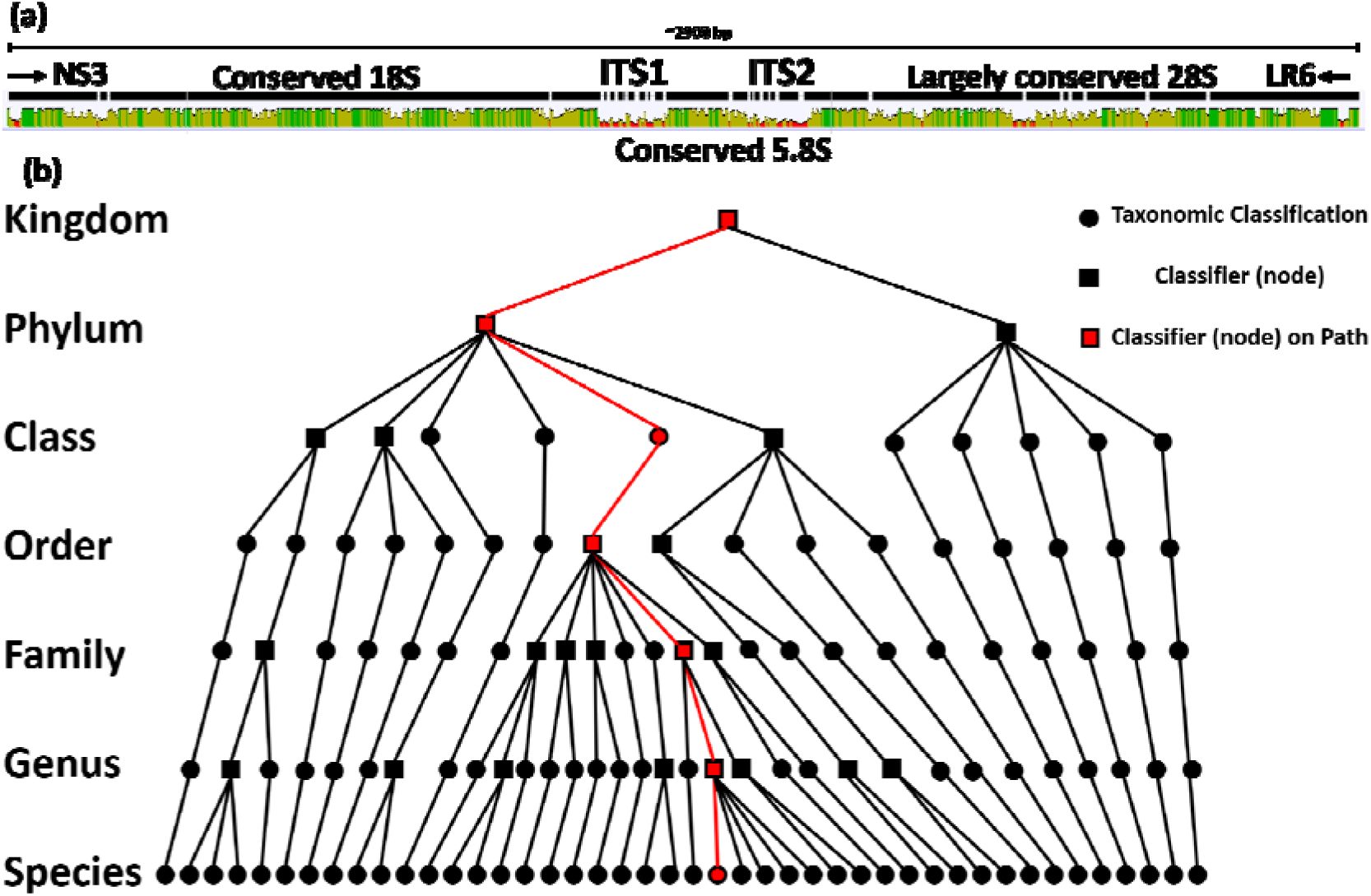
Visualisation of the fungal ribosomal DNA region and machine learning decision tree. **(a)** The fungal ribosomal DNA region between the NS3 and LR6 primers covers around ∼2900 bp. Shown is the alignment of consensus sequences (created with Geneious Prime), with highly conserved regions in green and variable regions in red. **(b)** Taxonomic information for 44 species was used to create a decision tree showing the relationship between samples at each taxonomic rank. Where two or more species shared a common taxon, a machine learning classifier (node) was created to distinguish between those taxa. These classifiers were chained together to create a decision tree, whereby the classification of a read was undertaken by a cascade of classifiers. An example path down the decision tree is shown for *Candida albicans*.

With the advent and increasing use of long-read sequencing, some of the limitations of short-reads can be bypassed [8]. With long-reads, an extended ITS region can be sequenced encompassing ITS1 and ITS2 in addition to the minor variable regions of the 18S and 28S rRNA subunits using a single ‘universal’ primer set (Figure 1a) [9–12]. Long-read sequencing technologies from Pacific Biosciences (PacBio) have been extensively benchmarked for fungal metabarcode sequencing [13]. Previous work by Tedersoo et al. [14] using the PacBio RSII and Sequel instruments suggest long-read sequencing offers higher resolution for species identification than short-reads. Specifically, they found that using full-length ITS sequences increased identification precision for fungi by 33% compared to using ITS1 or ITS2 alone. Further, PacBio long-reads better reflected the community composition of a template mock community than traditional Illumina MiSeq short-reads [13, 14]. In contrast, Oxford Nanopore Technologies (ONT) products have not been as extensively benchmarked for fungal species identification using metabarcodes [15]. This is despite the fact that the ONT MinION has the unique advantage of being highly portable for field studies.

Here, we apply ONT long-read sequencing to fungal ribosomal DNA amplicons generated with the NS3 and LR6 primers [16], spanning close to 2900 bp in size (Figure 1a). We refer to this amplicon hereafter as the fungal ribosomal DNA region. Nanopore sequencing introduces a relatively high read error of around 10% at the time of conducting our study [15], but this read error has recently been improved to ∼5% with the ONT super high accuracy basecalling models. These high error rates make individual reads less suited for species identification using DNA metabarcodes with currently existing sequence clustering, alignment and k-mer based methods because the genetic distance of the variable regions between closely related species are often lower than the per read error rate [17]. In addition, the entries in most fungal DNA barcode databases, such as the NCBI ITS database and UNITE [18] are relatively short, with a median sequence length of 580 bp and 540 bp, respectively. This limits the analysis capacity of long-reads which encompass both ITS sequences and include minor variable regions in both 18S and 28S rRNA (Figure 1a).

The fungal kingdom is diverse, with an estimated 1.5-5 million species globally, performing important ecosystem functions [19]. For example, mycorrhizal fungi play a key role in regulating the nutrient uptake of almost all land plant species [20] and can improve plant growth under adverse conditions [21], while detritus-degrading saprotrophic fungi play an important role in improving soil fertility [22]. However, fungal pathogens can have severe adverse effects on human and animal health and agriculture. Fungi can cause large-scale biodiversity losses [23, 24], as demonstrated by the near extinction of amphibian taxa by the globally devastating fungal pathogen *Batrachochytrium dendrobatidis* [25], and the local extinction of several myrtaceae tree species by the rust fungus *Austropuccinia psidii* [26]. Fungal plant pathogens cost billions of dollars in global food production annually [27].

The importance of fungi warrants the development of improved sequence-based detection methods for fungi. In our current study we address the shortcomings of current techniques and assess the applicability of novel machine learning-based sequence analysis methods for long-read metabarcodes from the fungal kingdom. We focus on the detection of known knowns – taxa known to be present in a system – in the context of pathogen infections or host-associated samples. We explored the potential of machine learning classifiers as an alternative method for assigning individual error-prone long-reads to taxa, especially for detecting known or likely pathogen candidates. We demonstrate that our approach works on real data, and demonstrate that machine learning has a place in the pathogen detection space as a supplementary tool with specific functions, rather than as a replacement for existing methods.

Within the existing machine learning methods for identifying patterns across various data types [28–30], convolutional neural networks (CNNs) are one method especially suited for identifying the deterministic spatial relationships in a DNA sequence [31–34]. We applied a CNN approach to metabarcoding-based fungal species identification using a uniquely labelled sequencing dataset of a 2900 bp fungal ribosomal DNA region from 44 individually sequenced fungal taxa. This included species and strains associated with plants, from environmental and soil samples, and human pathogens. We used real-world datasets to closely mimic real-world sequencing conditions and demonstrate the potential of machine-learning in the pathogen detection space, and further demonstrate that this method can be used alongside simulated read datasets. We compared our machine learning approach to three commonly used analysis approaches, including alignment and k-mer based methods, on different in house and publicly available databases. We demonstrated its potential advantage on classifying closely related fungal species with few basepairs difference within the fungal ribosomal DNA region. Furthermore, by including a broad list of fungal taxa across the fungal kingdom and a comprehensive and focused group of fungal taxa related to the target species when training the machine learning models, the approach can identify and detect a known or suspected target species from a complex sample even if not all species present in the sample were included in the training dataset.

## RESULTS

### Design of a decision tree for machine learning classifiers for taxonomic assignment of fungal species

Here we explored the application of machine learning on individual nanopore reads for fungal taxonomic classification. We sequenced the fungal ribosomal DNA region of 44 fungal taxa individually to generate a labelled real-world dataset for which the ground truth is known for each individual read. This makes our dataset uniquely suited for our supervised machine learning approach and for benchmarking studies when comparing this to commonly used classification approaches. Our fungal species dataset included 39 ascomycetous taxa, spanning 19 families and 27 genera, in addition to five basidiomycetes. Given the high sequencing depth of the amplicon sequencing, we performed stringent quality-control steps on all reads to eliminate sequencing biases. We first filtered reads based on homology against a custom-curated database of the fungal ribosomal DNA region, to remove partial reads or reads from other areas of the fungal genome with partial primer binding. We then filtered reads by length to remove reads that were not within a 90% confidence interval around the mean read length of the fungal ribosomal DNA region for each taxon (see Supporting Information Table T1). The *Geotrichum candidum* (former *Galactomyces geotrichum*) sample had too few reads for further processing, hence we complimented those with simulated reads using NanoSim [35]. This resulted in an average of 54,832 ± 35,537 reads available across all taxa. We took a subsample of these quality-controlled reads and split them into a training set and a test set, containing 85% and 15% of the subsampled reads respectively, to be used for training the machine learning classifiers and assessing the performance of the newly generated machine learning classifiers, respectively. We implemented a decision tree to be able to classify individual reads at each taxonomic rank from phylum to species (Figure 1). The taxonomic information for the 44 available individually sequenced taxa was used to create the cladogram for this decision tree. For simplification, we counted individual strains as separate species for this purpose to test the ability to discern between very closely related taxa. We generated one machine learning classifier for each node in our decision tree, where two or more sub-taxa diverged along the cladogram (Figure 1).

For training each of these classifiers, a balanced dataset was used, such that each possible outcome of the machine learning classifier had an equal number of reads. By giving equal weight to each group within the classifiers, we ensure that the machine learning model can identify the best distinguishing features between groups. These individual classifiers at different taxonomic ranks had a mean recall rate of 97.9 ± 1.1% for correctly classifying reads using the test read dataset. The lowest recall rate belonged to the species-level classifier that distinguished between *Candida* species, with a recall rate of 94.4%. The receiver operator characteristic (ROC) curves demonstrated a good classifier (see the GitHub repository at https://github.com/teenjes/fungal_ML).

To fully classify a read, we used the cladogram as a decision tree to link individual machine learning classifiers at each taxonomic rank. This allowed us to chain classifiers together to classify a read at each taxonomic rank, moving through the tree from phylum to species assignments. The outcome of a classifier at one taxonomic rank was used to decide the path along the tree, and thus this decision defined which classifier was appropriate for use at the next lower taxonomic rank (Figure 1). We refer to a classifier by the taxonomic rank that it outputs. For example, a species-level classifier takes reads from a specific genus and outputs a species, while a class-level classifier takes reads from a specific phylum and outputs a decision on the taxonomic class of the read. The recall rate of the individual classifiers at different taxonomic ranks can affect the final species-level recall rate for each individual read as it moves through the decision tree. This means that the final species-level recall rate is equal to or worse than the individual species-level classifier’s recall rate. Since not every path through the decision tree had a node at each taxonomic rank, the total number of nodes with classifiers is less than the total number of taxa. For example, the basidiomycete species *Puccinia striiformis* f. sp. *tritici* has only two classifiers, at the phylum level and the class level. The latter decides the class classification, but as *P. tritici* is the only species in the class Pucciniomycetes in our sequencing dataset, no further nodes were required to distinguish between this species and other taxa present in our dataset. Further, both the *Discula quercina* and *Tapesia yallundae* species were represented in two samples with different strains, although each was classified as a species for the purposes of this model. In total we trained 22 classifiers to distinguish our 44 fungal taxa.

### Machine learning methods provide robust classification, especially at the genus and species level

We compared the machine learning decision tree to other, more standard methods for read classification to determine the effectiveness of this technique. We assessed the ability of the other methods at classifying reads across multiple taxonomic ranks because the tiered nature of the decision tree offers the potential to gleam taxonomic information from a read, even when it cannot be confidently classified at the species level. We first looked at the use of a naïve Bayes classifier, however this performed very poorly and was inconsistent in the results produced (data not shown), which is consistent with previous studies [34]. We used two additional classification techniques. We applied *minimap2*, a pairwise alignment-based method designed to be used with long-reads, against a gold-standard custom-curated database generated from the consensus sequences of all 44 species present in the decision tree (gold standard alignment). This is the most appropriate comparison for our machine learning approach because the gold standard and machine learning approaches are directly derived from our sequencing dataset. To compare the machine learning approach with methods where the sequencing data was not used to create the classification database in some way, we applied *minimap2* to a large publicly-available database of fungal ITS sequences from NCBI [36, 37] (NCBI alignment), and applied *Kraken2*, a k-mer-based classifier, to the same NCBI database (*Kraken2*).

To compare these methods, an *in silico* mock community was generated from our labelled sequencing data via subsampling for which we know the ground truth classification for each sequencing read. We selected 13 species from the original 44 taxa, including some that required four or more classifiers to generate the machine learning decision tree, particularly those from populous genera with closely related species. Although all species from this mock community were present in the gold standard database, the NCBI database was missing some genera and species. All of these missing or unclassified taxonomies were recorded as having a recall rate of zero percent, artificially decreasing the quality at lower taxonomic ranks.

Our machine learning decision tree approach maintained a consistently high recall rate across all taxonomic ranks, with a mean species level recall rate of 93.0 ± 2.8% (1σ). Notably, it performed very well for closely related taxa, including the cryptic species *Candida metapsilosis* and *Candida orthopsilosis* and another closely related species *Candida albicans*. The two cryptic *Candida* species (*C. metapsilosis* and *C. orthopsilosis*) had a very high consensus sequence similarity, with a genetic distance of 2.74% (97.26% identity) in our fungal ribosomal DNA target region representing the genetically least distinct species pair. Our machine learning approach achieved species level recall rates of 90.1% and 89.1% for *C. metapsilosis* and *C. parapsilosis*, respectively, even with per read error rates of about 10%. This highlights the strength of our approach.

The gold standard alignment approach also performed very well when compared to the machine learning approach across all taxonomic ranks (Figure 2). The majority of the species were classified with recall rates in excess of 95%. Yet this approach significantly underperformed when trying to differentiate taxa with low genetic distance such as those from the *Candida* genus. As with the machine learning approach, the three *Candida* species were classified with the lowest recall rate at the species level, with *C. albicans, C. metapsilosis* and *C. parapsilosis* being classified with recall rates of 35.8%, 34.0% and 57.5% respectively. These difficulties are also reflected in the overall mean species level recall rate of 76.6 ± 25.5%, which is much lower than our machine learning approach.

**Figure 2:**
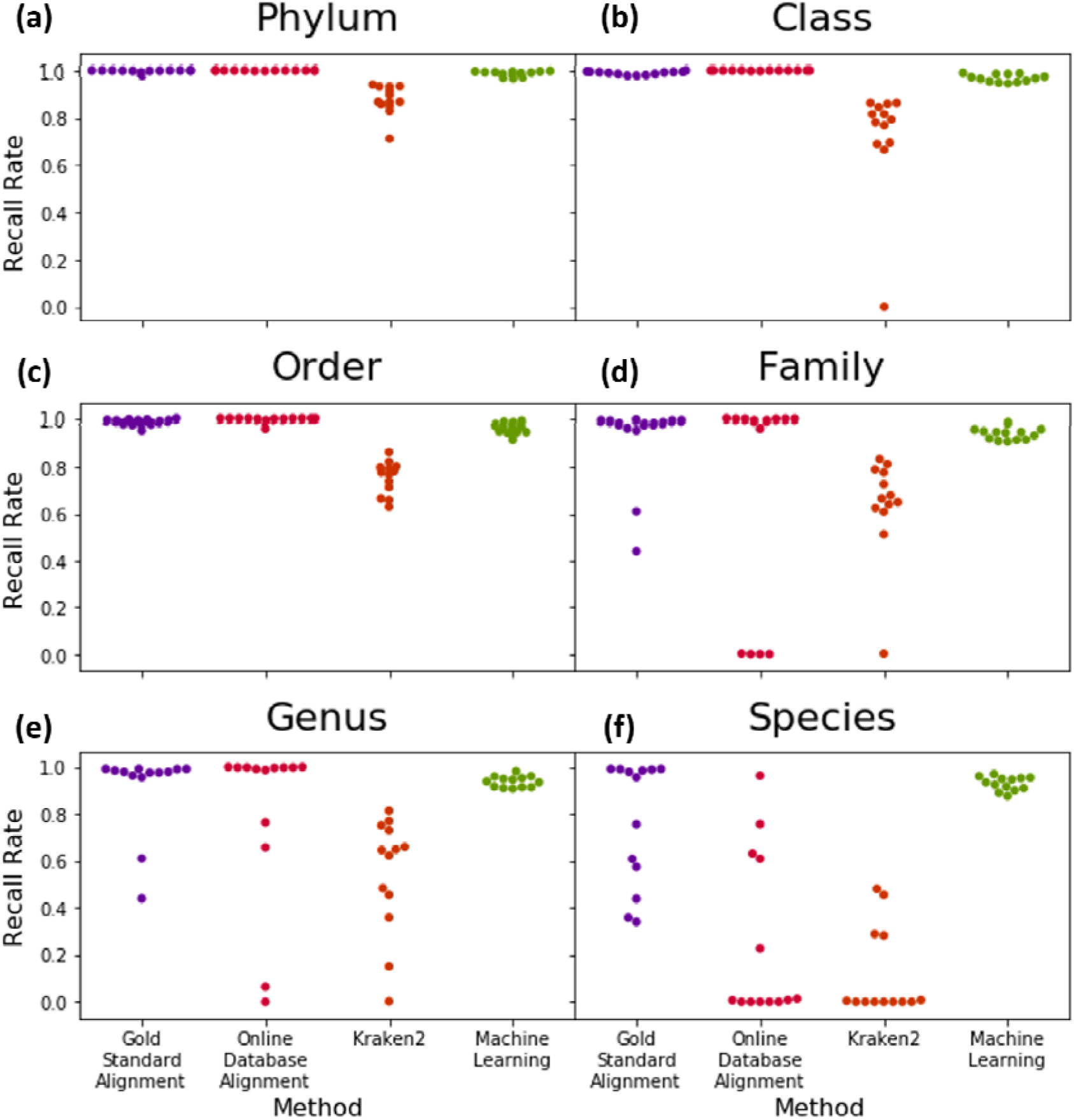
Machine learning based species identification performs especially well for closely related species. Recall rates of the alignment-based *minimap2* technique, k-mer-based *Kraken2* method and the machine learning decision tree across different taxonomic ranks **(a-f)**. The *minimap2* technique, as applied to the gold standard database, was successful across most taxonomic ranks, but lower recall rates were recorded for closely related species at the species level **(f)**. Both the *minimap2* and *Kraken2* methods were applied to the NCBI database, and while the *minimap2* NCBI alignment was more accurate across most taxonomic ranks, both showed comparable recall at the species level. The machine learning decision tree approach provided the greatest classification power for closely related species, despite lower recall rates for some distantly-related species than the gold standard alignment method.

Next, we assessed our dataset with alignment and k-mer based analysis approaches when using the publicly available NCBI database. Overall, NCBI alignment with *minimap2* performed similarly well at higher taxonomic ranks. However, inconsistent or missing naming conventions at the family level and missing or alternate species labels, meant that the overall recall rate was low at the species level, although most of the samples were classified with a high recall rate at the genus level. This low species level recall rate is an artefact created from the choice of database, which is reflected in the similarly poor species level recall rates of the *Kraken2* method. Overall, the k-mer based *Kraken2* was less accurate than all other methods tested across all taxonomic ranks.

### Identifying target species from a complex sample of unknown composition using the machine learning decision tree

A key feature of a species classification tool is its ability to identify a known target species from a complex sample of unknown composition. This is especially important when attempting to identify the presence of a target species, such as a specific pathogen, from a patient or environmental sample.

We generated two additional sequencing datasets of unknown composition from two distinct environments to test the capability of our machine learning decision tree to identify a given target species. These datasets were generated with the same PCR and sequencing protocols as for the individual 44 training species focusing on the fungal ribosomal DNA region. The first dataset was derived from amplicon sequencing of fungi-infected wheat leaves (wheat dataset) and the second was derived from bronchoalveolar wash in a clinical setting (clinical dataset). To each of these sequencing datasets of unknown composition, we spiked *in silico* a known number of reads with known labels as test case. We choose *Aspergillus flavus*, a crop pathogen, and *Candida albicans*, a human pathogen, to compare two species without and with very close sister species in our linked machine learning classifiers. We then tested recall and false positive rate of our machine learning classifiers using our *in silico* spiked reads, assuming that the original datasets of unknown composition did not contain any reads of either species.

We first plotted the propagated confidence score of the species level classification for all reads in each *in silico* spiked dataset to better understand the behaviour of our machine learning decision tree on samples containing reads of unknown origin (Supporting Information Figure S1). This clearly shows that the propagated confidence scores for reads of unknown origin are far lower than reads of species the classifiers were trained on. We then assessed the recall and false positive rate of the *in silico* spiked datasets at different confidence scores thresholds assuming that the original sample did not include any reads of the *in silico* spiked in species (Figure 3). Increasing the thresholds reduced the recall and false positive rate in both cases. For *A. flavus*, the recall rate remained above 90% until the confidence threshold reached 0.9, and the false positive rate was consistently low across both the clinical and wheat datasets with reads of unknown origin. A confidence threshold of 0.85 resulted in a high recall rate of 0.917, while maintaining a low false positive rate of just one percent. For *C. albicans*, not using a confidence threshold at all resulted in a recall rate of 87.7% and false positive rate of 11.7%. However, by using a confidence threshold of 0.85, the recall rate was only decreased to 72.4% while reducing the false positive rate to only 1.7% in the clinical dataset. We recommend this confidence score threshold of 0.85 as suitable for retaining a high recall rate while achieving a low false positive rate, even for a member of a difficult-to-distinguish genus like *Candida*.

**Figure 3:**
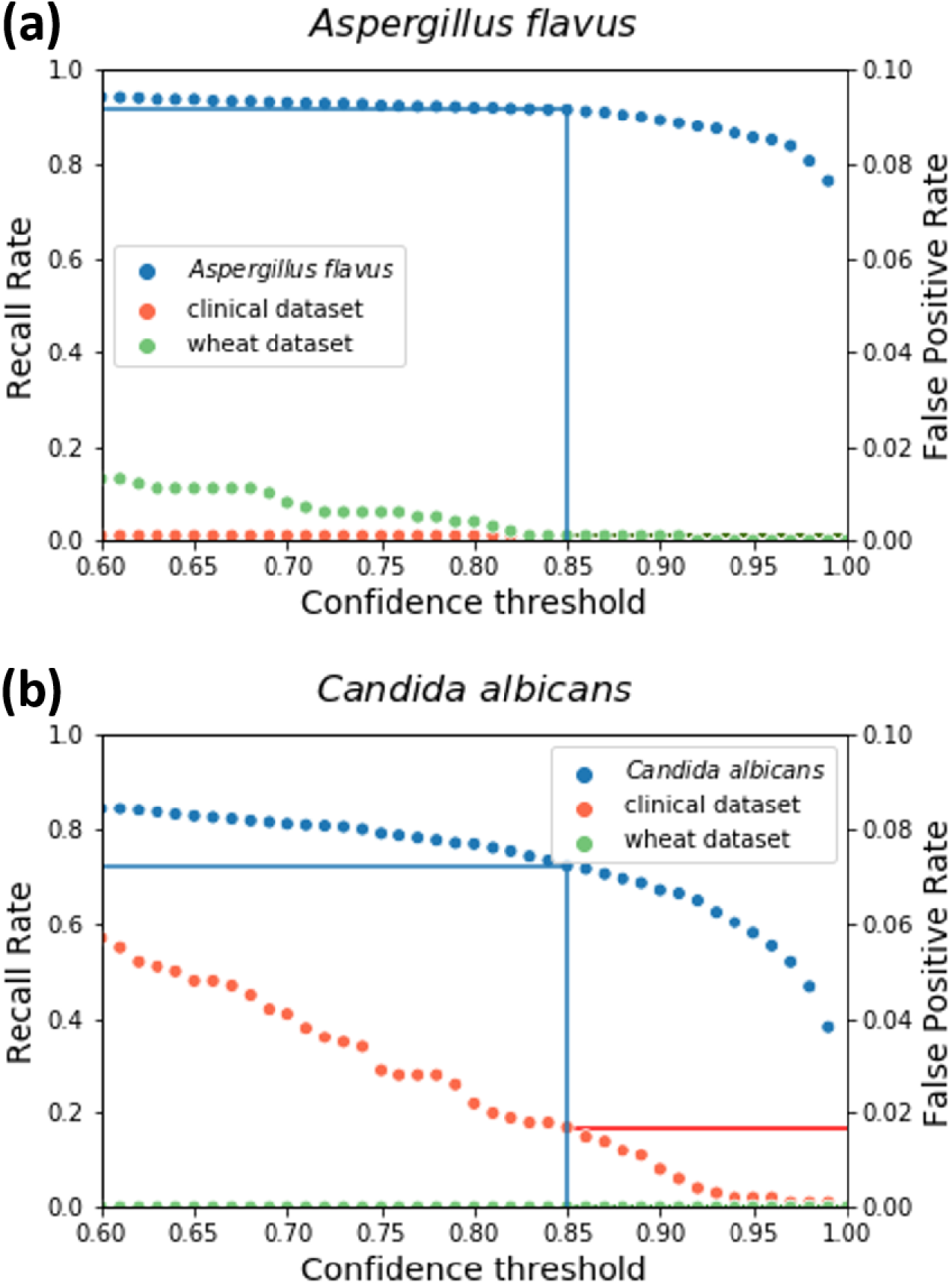
Varying the confidence score threshold affects the recall rate and false positive rate when identifying target species from complex samples of unknown composition. The plots show the recall rate (left axis, blue dots) and false positive rate (right axis, orange or green dots) at varying propagated confidence score thresholds for *Aspergillus flavus* **(a)** and *Candida albicans* **(b)** when spiked into clinical (orange) or wheat (green) datasets. Both plots are based on 2000-read *in silico* spiked samples containing 1000 reads with known labels (*A. flavus* or *C. albicans*) and 1000 reads of unknown origin. For *A. flavus*, a confidence threshold of 0.85 maintains a recall rate of 91.7%, while reducing the false positive rate to 1% for both datasets. For *C. albicans*, the same confidence threshold of 0.85 has a recall rate of 72.4% and reduces the false positive rate below 2% for both datasets.

### Training a machine learning classifier with simulated reads resulted in similar accuracy to those trained on real data

Previously, models were trained using sequence data from pure fungal culture. However, this is not always accessible, and limits the ability for the approach to be extended. This allows for samples with low read counts, such as the *G. candidum* sample, to be augmented to have sufficient read counts for training and enables rapid extension of the suggested approach to more species. We simulated 35,000 reads for *A. flavus* based on our reference ribosomal DNA sequence using NanoSim v3.0.0 to test whether training machine learning models using simulated reads would produce the same level of accuracy as training with reads generated by sequencing amplicons generated from fungal reference cultures.

We retrained all models in the decision tree path of *A. flavus* using these simulated reads. None of the resultant models were statistically different to those models trained using real sequenced data in terms of precision or recall. Further, they showed similar ROC curves (see the GitHub repository at https://github.com/teenjes/fungal_ML). This suggests machine learning classifiers can be trained using simulated reads and still accurately distinguish between taxa across all taxonomic ranks.

We also used these modes trained using simulated data to test whether the use of simulated training data affected the ability of the system to detect target species from a complex sample. We found no difference in the recall or false positive rates for detecting the target species using these simulated-trained models, with a near identical distribution to that shown in Supporting Information Figure S1, and a recall rate of 0.910 when using a confidence threshold of 0.85 compared to the recall rate of 0.917 when trained on real data.

## DISCUSSION

Nanopore sequencing offers portable sequencing platforms and generates long-reads that can cover extended metabarcodes that include more sequence information than the classic metabarcodes from Illumina short-read sequencing [38]. The use of long-read nanopore sequencing is increasing, and analysis pipelines have been developed for real-time species identification on the MinION, such as the ‘What’s in my pot’ pipeline for complex sample analysis [39]. Yet currently, metabarcode datasets in publicly available databases are limited in barcode length and often do not cover these extended regions, limited only to ITS1 and ITS2 regions of ∼580 bps. This poses challenges when classifying error-prone long-reads into different species when the genetic differences within those metabarcode regions are less than the differences from sequencing errors [40].

Here, we implement a novel machine learning approach for species level classification. We used a tree descent approach, with a similar strategy to the IDTAXA taxonomic classifier [41], although the objectives differ. IDTAXA also uses a tree-like approach to classify a subsample of the k-mers into a given taxonomic rank using machine learning, and the user can specify certain confidence thresholds to avoid over-classification when confidence is low [41]. While IDTAXA’s goal is to classify unknown reads to a confident taxonomic rank, our approach is more focused on the detection of one or a few target species from mixed samples. As such, we used full-length reads rather than k-mers from the short ITS regions, as used by Vu et al. [34], for training the machine learning models. Training these models with long reads is computationally less efficient than with k-mers, but is more suitable for detection-focused objectives. The use of only the ITS regions themselves, rather than the extended read used here, may be feasible for detection-focused tasks, but may lose resolution gained from small variations in the otherwise conserved RNA-coding regions that surround the ITS. As such, very closely related taxa may be more difficult to distinguish if only the non-coding ITS is used [34].

Our approach showed great potential on improving the accuracy of sequencing-based fungal classification at a species level. For example, some closely related species, such as those from the *Candida* genus, were readily misclassified with a recall rate lower than 50% using the gold standard alignment strategy. This is indicative of the problems of alignment-based classification methods for fungi, especially given the relatively high error rate of the nanopore long-reads [42]. Using our machine learning method, the same group of species was classified with recall rates of at least 90% (Figure 2f). This is remarkable given the per-read error rate of 10% for the nanopore reads used, which is much larger than the genetic distance of 2.74% observed between some closely related taxa. With recent improvements in the sequencing and basecalling of nanopore reads reducing the per-read error rate [43], the resolution of our method is likely to increase.

To ensure accurate classification of reads from closely related taxa using our machine learning classifiers, the training of species-level classifiers must include metabarcodes from other taxa closely related to the target for detection. If closely related taxa are not included in training, the likelihood of false positive hits increases as closely related taxa may be identified as the target taxon even when the target taxon is not present. For example, *Candida metapsilosis* may be falsely identified as *Candida albicans* if the model was not trained on both species. As such, accurate species-level identification requires that machine learning classifiers are trained on long metabarcodes from as many fungal species in the same genus as possible. This is especially important when a genus contains both pathogenic and non-pathogenic species. In this way, our approach might be particularly applicable to targeted diagnostic tasks in specific settings, such as detecting fungal pathogens in agriculture [44], forest management [45] and medicine [46], or screening imports for invasive pathogens in aid of border biosecurity [47, 48].

The initial comparisons were based on idealised databases derived from our sequencing dataset, for which metabarcode length and online database entry length were equivalent. Hence, is it expected that analyses based on these idealised databases will outperform other approaches relying on public databases with short reference sequences of uneven length, as shown by our results (Figure 2). Interestingly, the *Kraken2* approach underperformed compared to the alignment-based approach in our current study. This is consistent with previous studies with nanopore data, where the classification accuracy of *Kraken2* never exceeded that for BLAST, another alignment-based classification program, when using the default 35 bp k-mers [49, 50]. One approach to improve the k-mer based classification accuracy for error-prone reads would be to use a smaller k-mer length. Another common issue of public fungal databases was that many species were not included in the database or were presented with different taxonomic labels, which resulted 0% recall rates of some families and species. Changing fungal nomenclature over time can be an issue when using these online databases for classifying fungal metabarcode reads, as the nomenclature is not always updated, leading to outdated or uncorrected taxonomic information persisting in databases [51, 52].

We also tested if our machine learning approach could accurately identify target species in complex samples of unknown composition without having classifiers for all fungal species present in the sample. These complex samples had previously been explored through shogun sequencing [53, 54]. We were able to show that by only training a limited set of classifiers we can detect target species with relatively low false positive and high recall rates in *in silico* spiked datasets with known ground truth of the spiked reads only. By adjusting the confidence score one can decide how much false positive and false negatives one is willing to tolerate. We found a threshold of 0.85 on the propagated confidence score at the species level classification was sufficient to reduce the false positive rate while maintaining high recall rates. Here, the false positive rate was misleadingly high for *Candida* species in the clinical sample and for *Aspergillus* species in the wheat sample. We naively assumed these species were not present in our complex sample save for those reads that had been spiked in. *Candida* species, including *C. albicans*, are common in humans, and similarly *A. flavus* is a common fungus present on wheat. However, it also demonstrates the need to curate closely related species for training the models, as other species from the *Candida* or *Aspergillus* genera may have been misidentified as the target species from these complex samples.

Further, we found that training the machine learning classifiers using simulated reads, rather than sequenced reads, did not disadvantage the model’s ability to distinguish between taxa at any level. It is useful for extending this approach to a larger training set with an increased number of species, particularly for including additional taxa closely related to the target species. We can simulate reads based on publicly available reference sequences to train the classifiers, making this approach accessible even where reference cultures are not readily available.

The taxa used to train the classifiers are flexible and can be changed to suit the user’s need including simulation based on publicly deposited genomes of unavailable species. For example, additional species from a specific taxon could be added to reduce false positive detections within that taxa. While many existing classifiers are designed for species classification, and can falsely identify previously unknown species as existing species by over-labelling [55], our approach focused on detection of known target fungi rather than identifying all species present. While unknown taxa absent in the training dataset will be assigned a label, the use of a confidence score threshold can aid in filtering these misclassified unknowns out. The principles behind the application of machine learning to the fungal ribosomal DNA region can be expanded to other barcoding regions, such as *cytochrome c oxidase I* [56] or *elongation factor 1 alpha* [46]. Recent developments by Oxford Nanopore Technologies to improve barcoding accuracy, cost-effectiveness and scalability, such as the recently-released Q20+ chemistry, offers promise for expanding our proof-of-concept study to more species using multiple barcodes to improve the species-level resolution and overall classification accuracy [57].

## CONCLUSIONS

Long-read sequencing can be more accurate for species identification and detection than short reads, offering an advantage for future use [13, 14]. Here, we propose a tangible solution for fungal classification and detection by applying a novel neural network-based machine learning approach on extended fungal ribosomal DNA barcodes using nanopore long-read sequencing. Our machine learning approach can identify target taxa with high accuracy from complex samples of unknown origin, making it applicable to pathogen identification in agriculture, biosecurity, and clinical settings. Our approach performs especially well on closely related species, outcompeting current alignment-based or k-mer-based classification methods for long reads. Future optimisations for real life applications are required in terms of computational efficiency and size of reference training sets.

## METHODS

### Fungal pathogen sample collection, DNA extraction and ITS amplification

We collected different fungal tissue differently for DNA extractions. The tissue collection processes for each fungal taxon are summarized in Supporting Information Table T1.

We used three different DNA extraction methods for all the species in the mock communities. The methods for each species are listed in the Supporting Information Table T1. Collectively, we used two commercially available kits: the Qiagen DNeasy Plant Mini Kit (cat. no. 69106) for most plant pathogenic fungi, and the Quick-DNA Fungal/Bacterial Miniprep Kit (cat. no. D6005, Zymo Research) for some human pathogenic fungi following the manufacturer’s protocol. We used a phenol chloroform-based DNA extraction method for other human pathogenic and environmental fungi modified from Ferrer et al [58]. Briefly, 100 mg of leaf tissue was homogenized and cells were lysed using cetyl trimethylammonium bromide (CTAB, Sigma-Aldrich) buffer (added RNAse T1, Thermo Fisher, 1,000 units per 1750 μl), followed by a phenol/chloroform/isoamyl alcohol (25:24:1, Sigma-Aldrich) extraction to remove protein and lipids. DNA was precipitated with 700 μl of isopropanol, washed with 1 ml of 70% ethanol, dried for 5 min at room temperature, and resuspended in 50 μl TE buffer containing 10 mM Tris and 1 mM EDTA at pH 8. For the human clinical sample and the field infected wheat sample, we directly used the DNA described in the original article [53, 54] for PCR amplification. Quality and average size of genomic DNA was visualized by gel electrophoresis with a 1% agarose gel for 1 h at 100 volts. DNA was quantified by NanoDrop and Qubit (Life Technologies) according to the manufacturer’s protocol.

We used the NS3 (5’ GCAAGTCTGGTGCCAGCAGCC ‘3) and LR6 (5’ CGCCAGTTCTGCTTACC ‘3) primers [16] to generate the fungal ribosomal DNA fragment of all samples, and the EF1-983F (5’ GCYCCYGGHCAYCGTGAYTTYAT ‘3) and EF1-2218R (5’ ATGACACCRACRGCRACRGTYTG ‘3) primers [16] were used to sequence a secondary region, the fungal elongation factor 1 alpha region, which was not used for assessing the machine learning method. We used the New England Biolabs Q5 High-Fidelity DNA polymerase (NEB #M0515) for the PCR reaction following the manufacturer’s protocol. Around 10 – 30 nanograms of DNA were used in each PCR reaction. After PCR, DNA was purified with one volume of Agencourt AMPure XP beads (cat. No. A63881, Beckman Coulter) according to the manufacturer’s protocol and stored at 4°C.

### Library preparation and DNA sequencing using the MinION

DNA sequencing libraries were prepared using Ligation Sequencing 1D SQK-LSK108 and Native Barcoding Expansion (PCR-free) EXP-NBD103 Kits from ONT, as adapted by Hu and Schwessinger [59] from the manufacturer’s instructions with the omission of DNA fragmentation and DNA repair. DNA was cleaned up using a 1x volume of Agencourt AMPure XP beads (cat. No. A63881, Beckman Coulter), incubated at room temperature with gentle mixing for 5 mins, washed twice with 200 μl fresh 70% ethanol, the pellet allowed to dry for 2 mins, and DNA eluted in 51 μl nuclease free water and quantified using NanoDrop^®^ (Thermo Fisher Scientific, USA) and Promega Quantus^™^ Fluorometer (cat. No. E6150, Promega, USA) following the manufacturer’s instructions. All DNA samples showed absorbance ratio A260/A280 > 1.8 and A260/A230 > 2.0 from the NanoDrop^®^. DNA was end-repaired using NEBNext Ultra II End-Repair/ dA-tailing Module (cat. No. E7546, New England Biolabs (NEB), USA) by adding 7 μl Ultra II End-Prep buffer, 3 μl Ultra II End-Prep enzyme mix. The mixture was incubated at 20°C for 10 mins and 65°C for 10 mins. A 1x volume (60 μl) Agencourt AMPure XP clean-up was performed, and the DNA was eluted in 31 μl nuclease free water. Barcoding reaction was performed by adding 2 μl of each native barcode and 20 μl NEB Blunt/TA Master Mix (cat. No. M0367, New England Biolabs (NEB), USA) into 18 μl DNA, mixing gently and incubating at room temperature for 10 mins. A 1x volume (40 μl) Agencourt AMPure XP clean-up was then performed, and the DNA was eluted in 15 μl nuclease free water. Ligation was then performed by adding 20 μl Barcode Adapter Mix (EXP-NBD103 Native Barcoding Expansion Kit, ONT, UK), 20 μl NEBNext Quick Ligation Reaction Buffer, and Quick T4 DNA Ligase (cat. No. E6056, New England Biolabs (NEB), USA) to the 50 μl pooled equimolar barcoded DNA, mixing gently and incubating at room temperature for 10 mins. The adapter-ligated DNA was cleaned-up by adding a 0.4x volume (40 μl) of Agencourt AMPure XP beads, incubating for 5 mins at room temperature and resuspending the pellet twice in 140 μl ABB provided in the SQK-LSK108 kit. The purified-ligated DNA was resuspended by adding 15 μl ELB provided in the SQK-LSK108 (ONT, UK) kit and resuspending the beads. The beads were pelleted again, and the supernatant transferred to a new 0.5 ml DNA LoBind tube (cat. No. 0030122348, Eppendorf, Germany).

In total, four independent sequencing reactions were performed on a MinION flow cell (R9.4, ONT) connected to a MK1B device (ONT) operated by the MinKNOW software (version 2.0.2): 11 species for each flowcell. Each flow cell was primed with 1 ml of priming buffer comprising 480 μl Running Buffer Fuel Mix (RBF, ONT) and 520 μl nuclease free water. 12 μl of amplicon library was added to a loading mix including 35 μl RBF, 25.5 μl Library Loading beads (ONT library loading bead kit EXP-LLB001, batch number EB01.10.0012) and 2.5 μl water with a final volume of 75 μl and then added to the flow cell via the SpotON sample port. The “NC_48Hr_sequencing_FLOMIN106_SQK-LSK108” protocol was executed through MinKNOW after loading the library and run for 48 h. Raw fast5 files were processed using Albacore 2.3.1 software (ONT) for basecalling, barcode de-multiplexing and quality filtering (Phred quality (Q) score of > 7) as per the manufacturer’s recommendations.

Raw unfiltered fastq files were uploaded into NCBI Short Reads Archive under BioProject PRJNA725648.

### Processing and manipulation of fungal pathogen reads

The read datasets obtained contained both the fungal ribosomal DNA sequence and that of elongation factor 1 alpha. A two-step data filtration process was applied to remove the elongation factor 1 alpha reads and outlier reads from the fungal ribosomal DNA read set.

Reads were mapped to an in-house database of fungal ribosomal DNA regions to select reads of a similar general structure to the ITS region. This homology-based filter assumes the structure of the fungal ribosomal DNA region is similar between species due to shared ancestry, repeatedly shown to be true [60]. The in-house database was curated from 28 ITS sequences from the NCBI Nucleotide database, from a range of genera across the fungal kingdom. This process mapped reads using *minimap2* (v2.17), using the map-ont flag. Reads that failed to map were discarded.

Remaining reads were filtered for read length. The expected read length for the fungal ribosomal DNA region varied by species, averaging 2600-3200 bp. As the mean length and spread of successfully filtered reads differed between samples, a 90% confidence interval cut-off around the mean read length was applied. This interval was sufficient to exclude short or very long reads resulting from incomplete or partial homology filtering or errors in the sequencing or basecalling processes.

### Augmenting read datasets

Some reads were simulated using a consensus sequence and MinION read error profile to ensure all samples had at least 15,000 reads for use in the design of the machine learning classifiers downstream. NanoSim (v2.0.0) [35] was used for one species, *Geotrichum candidum*, to generate an additional 8,782 simulated cDNA reads. These reads were generated using an identical error profile and length spread to the pre-existing non-simulated reads.

An additional simulated dataset was later generated using NanoSim v3.0.0 for use in testing the ability to train models using simulated data based on references available online, without losing the ability to accurately classify reads from real-world data. A total of 35,000 additional reads were simulated for *Aspergillus flavus* to ensure the maximum number of reads that could be processed using our approach was available. These reads were generated using an error profile identical to the pre-existing non-simulated reads but were based on a reference consensus sequence for *A. flavus*.

### Generating consensus sequences for each species

The consensus sequence, an aggregate sequence formed from the comparison of multiple sequences that represents the ‘true’ sequence, was generated using 200 randomly subsampled filtered reads for each sample. Primer sequences were removed using *Mothur* v1.44.11 [61], an alignment file was generated using *muscle* v3.8.1551 [62] and the consensus sequence was generated from this file using *EMBOSS cons* v6.6.0.0 [63]. These consensus sequences were used to confirm taxonomy when species was initially unclear by mapping to the UNITE, SILVA and NCBI databases.

### Determining the relationships between samples

Using the taxonomic information available for each sample in MycoBank and the results of a BLAST search with the generated consensus sequences, a cladogram was designed to show the relationships between samples based on major taxonomic ranks. A machine learning classifier would be required at each point where two or more samples split on the cladogram (a node) to distinguish between samples for each read, with the cladogram functioning as a decision tree with the node-situated classifiers determining the path of tree descent. Individual strains of the same species were counted as separate species for this cladogram.

### Creation of neural network classifiers to distinguish between samples

A convolutional neural network (CNN) was chosen as the most appropriate type of machine learning classifier due to its ability to use the spatial relationships between data features in the reads, such as the distance between ITS and other variable groups. CNNs are capable of learning from both minor variation and higher-order features, which is of particular importance given the high read error of nanopore reads. While we did compare a naïve Bayesian and a fully-connected multilayer perceptron model to the CNN approach, these produced quite poor results in comparison, and previous studies using short-read k-mers had shown CNNs to provide an advantage over the existing Bayesian Ribosomal Database Project and other machine learning classifier types [34]. Given this, we selected CNNs to be the basis of our decision tree moving forward.

CNNs work best when there is a balanced number of items in each classification class. As such, for each multiclass node on the cladogram, an equal number of reads were subsampled from each group of samples that would be represented in the node. So, for machine learning classifiers distinguishing between species, each species present contributes an equal number of reads, while at the kingdom level, each phylum contributes an equal number of reads, with said reads being distributed equally amongst all species belonging to that phylum. The number of reads subsampled was based on the largest number of reads available for each sample, with a maximum of 35,000 reads due to computational processing limitations. For each read subsampled, the nucleotide sequence was converted to a numeric sequence, where A, C, G, and T became 0, 1, 2, and 3, respectively. This integer encoding produced the same result as using one-hot encoding to multiple dimensions. As not all sequences were of equal length, but an equal length was required to avoid sequence length being a distinguishing factor in the classifier, all sequences were padded out to a length of 5,000 bp. The padding used a value of 4 to avoid the padding data from affecting the identification of key features for classification.

Each read was assigned a label representing the output class it would belong to in the one-hot format. Labelled reads were then separated into a training set and a test set. The training set contained 85% of the reads, and was used to train the machine learning classifiers, while the test set contained the remaining 15% of labelled reads and was used to test the efficacy of said classifiers on similar data that the classifier had not previously encountered. This was done using the leave out N sequences approach rather than leaving out N taxa, which is more appropriate to represent the ‘dark matter’ taxa, as we wanted to focus on detecting the known knowns rather than the potential unknowns [55]. The neural network was created using the Sequential classifier of the *Keras* framework for neural networks [64], containing five layers of neurons and trained across 10 epochs.

Specific details for the design of the machine learning classifiers and the required software packages for machine learning and other analyses can be found at https://github.com/teenjes/fungal_ML.

### Evaluation of the machine learning classifiers

The test set was used to assess the accuracy of the various machine learning classifiers. As the test set data was labelled, the expected outcome for each read was known, and could be compared to the output of the machine learning classifier. The accuracy, or classification rate, of these classifiers was the proportion of reads in the test set for whom the prediction of the machine learning classifier, as determined by the highest confidence score, matched the expected outcome. This is equivalent to the recall rate [1], where matches to the expected outcome were true positives and matches outside this outcome were false negatives.

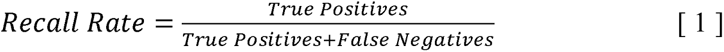

### Chaining machine learning classifiers into a decision tree

When seeking to identify members of a specific taxon in a community, where the members are not immediately obvious from the species name, it is useful to have samples classified at each taxonomic rank. A singular classifier would require excessive computational power to do this. As such, we chained the machine learning classifiers together into a decision tree with a tree descent approach, based on the cladogram of the species present in our sample. The most confident outcome of the machine learning classifier at one taxonomic rank would be used to decide the path along the decision tree. This path could either lead into another machine learning classifier, if the path diverged again, or lead all the way down to the species level with the same confidence.

### Alternative methods for fungal pathogen read classification

We used two different commonly used methods for fungal pathogen metabarcode classification to benchmark the linked machine learning model approach: an alignment-based method in *minimap2*; and a k-mer-based method in *Kraken2*. To compare these methods, we generated an *in silico* mock community from our labelled sequencing data for which we know the ground truth classification for each sequencing read. This mock community contained 13 of the 44 species used to generate the original machine learning decision tree, randomly subsampling 1000 reads from those not previously used for training the machine learning classifiers. Species were selected to focus on species for whom multiple machine learning classifiers would be required, in particular species with populous genera.

For this *minimap2*-based alignment method, two separate databases were used for identification. Firstly, a gold standard database was created in-house to represent the best-case scenario for identification, when all the species present in a sample are also present in the database. This contained the labelled consensus sequences of all 44 species present in the machine learning decision tree, using the consensus sequences already generated from 200 randomly selected filtered reads. The second was a publicly available database of fungal ITS sequences from NCBI (ftp://ftp.ncbi.nlm.nih.gov/refseq/TargetedLoci/Fungi/fungi.ITS.fna.gz, downloaded Feb 2021). *Minimap2* was applied to each of these databases using the map-ont flag. As the alignment tool can return multiple hits if alignment is good enough, only the best hit was taken for each read.

We used *Kraken2* (v2.0.8) to assign the NCBI taxonomic ID for the same 1000 reads of each species as used in the machine learning decision tree. We generated a *Kraken2* NCBI ITS database with the same fasta file downloaded from above. We used the *Kraken2*-build command with the --add-to-library and --build flag. We used the Python pandas module to modify the *Kraken2* output file and the numpy module to calculate the accuracy.

### Identifying a key species from a complex sample using machine learning

To assess the suitability of machine learning for this problem, we utilised the two complex datasets sampled from fungi-infected sources of unknown compositions: the field infected wheat dataset [53] and the human clinical dataset [54], to create *in silico* mock communities. We randomly subsampled 950 reads from these datasets, after initial processing as done on the data for machine learning, and spiked in 50 reads from one of two target species with known ground truth: *Aspergillus flavus*, a crop pathogen; and *Candida albicans*, an opportunistic human pathogen present in the human microbiome. This created a total of four 1000-read synthetic communities, two of which paired a target species and dataset from the same source (*A. flavus* with the wheat dataset and *C. albicans* with the clinical dataset) and two communities where the target species would not be expected to be present in the complex dataset unless it had been spiked in. We used the propagated confidence scores for assessing the recall rate for these spiked datasets, where the confidence score at each taxonomic rank was multiplied to give a final overall confidence at the species level.

We created an additional four *in silico* mock communities to assess the change in recall rate and false positive rate [2] as a confidence threshold was applied.

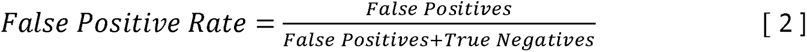

Each mock community was created by randomly subsampling 1000 reads from one of *A. flavus* or *C. albicans* samples with known ground truth and adding an additional 1000 randomly subsampled reads from one of the wheat or clinical datasets containing reads of unknown origin. In total, this resulted in four 2000-read *in silico* mock communities. We assumed the datasets with reads of unknown origin did not contain any reads for the target species tested, placing an upper bound on the false positive rate and a lower bound on the true positive rate. Any positive identifications of the target species *A. flavus* or *C. albicans* with a propagated confidence score below the confidence threshold were instead classified as negative identifications.

### Identifying a key species from a complex sample using classifiers trained on simulated reads

We used the augmented simulated read dataset from *Aspergillus flavus* generated by NanoSim to retrain those five models involved in the decision tree path leading to *A. flavus*, with an identical process to the initial model training. These models were incorporated into the decision tree in place of the original classifiers to test the ability of the decision tree to identify key species from complex samples when trained on simulated data. This used an identical method to the above method for identifying a key species from a complex sample.

## DECLARATIONS

### ETHICS APPROVAL AND CONSENT TO PARTICIPATE

Not Applicable.

### CONSENT FOR PUBLICATION

Not Applicable.

### AVAILABILITY OF DATA AND MATERIALS

The code generated and used for machine learning during the current study is available in the GitHub repository, available at https://github.com/teenjes/fungal_ML. The single-species datasets generated and/or analysed during the current study are available in SRA under BioProject PRJNA725648, available at http://www.ncbi.nlm.nih.gov/bioproject/725648. The complex mixed-species ‘clinical’ and ‘wheat’ datasets analysed during the current study are available in the same BioProject as SRX10704261 and SRX10704260, respectively.

### AUTHOR CONTRIBUTIONS

TE, YH, BM, and BS designed the experiments and performed the analysis. YH extracted fungal DNA and performed all sequencing reactions. LI, MTV, LMS, CCL, AWM, and WM provided fungal material and/or DNA. ES and JR provided feedback on experimental design and data analysis. WM, ES, JR, and BS provided funding for the project. TE, YH and BS wrote the manuscript. All authors commented on the manuscript and approved submission.

## ACKNOWLEDGEMENTS

We thank Jen Taylor, and Peter Solomon for suggestions on improving the machine learning models and determining statistics for identifying a species from a complex sample. We thank Peter Solomon and David Jones for providing fungal samples. This work was supported by computational resources provided by the Australian Government through the National Computational Infrastructure (NCI) under the ANU Merit Allocation Scheme.

