## Supplementary figures and images for "Linked machine learning classifiers improve species classification of fungi when using error-prone long-reads on extended metabarcodes"

### Supplemental Material

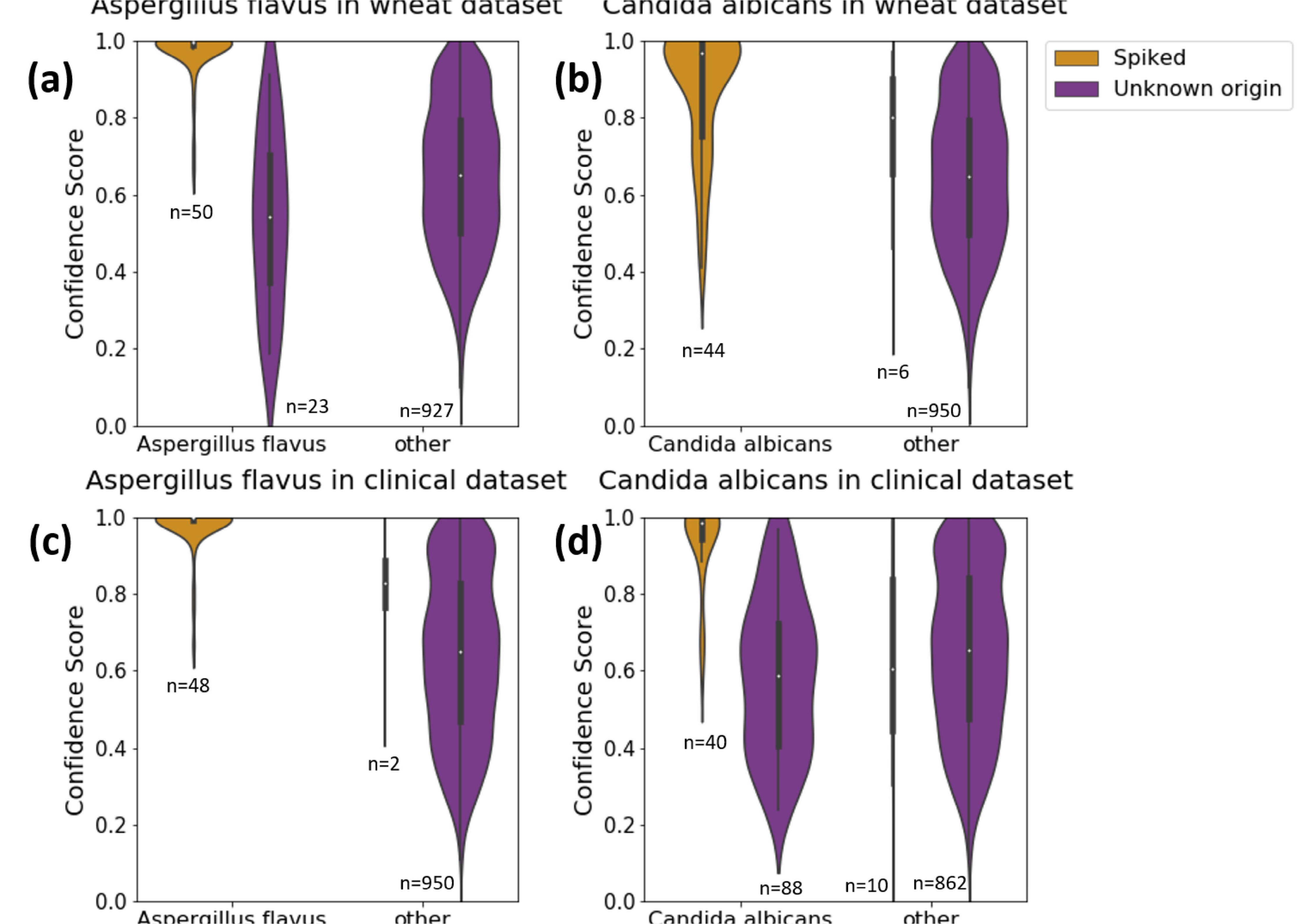
